# Alignment free identification of clones in B cell receptor repertoires

**DOI:** 10.1101/2020.03.30.017384

**Authors:** Ofir Lindenbaum, Nima Nouri, Yuval Kluger, Steven H. Kleinstein

**Affiliations:** Program in Applied Mathematics, Yale University, New Haven, CT 06511; Interdepartmental Program in Computational Biology and Bioinformatics, Yale University, New Haven, CT 06511; Department of Pathology, Yale University, New Haven, CT 06511; Department of Immunobiology, Yale University, New Haven, CT 06511; Center for Medical Informatics, Yale University, New Haven, CT 06511

## Abstract

Following pathogenic challenge, activated B cells rapidly expand and undergo somatic hypermutation, yielding groups of clonally related B-cells with diversified immunoglobulin receptors. Inference of clonal relationships based on the receptor sequence is an essential step in many adaptive immune receptor repertoire sequencing studies. These relationships are typically identified by a multi-step process that involves: (1) grouping sequences based on shared V and J gene assignments, and junction lengths, and (2) clustering these sequences using a junction-based distance. However, this approach is sensitive to the initial V(D)J gene assignments, which are error-prone, and fails to identify clonal relatives whose junction length has changed through accumulation of indels. Through defining a translation-invariant feature space in which we cluster the sequences, we develop an alignment-free clonal identification method that does not require gene assignments and is not restricted to a fixed junction length. This alignment-free approach has higher sensitivity compared to a typical junction-based distance method without loss of specificity and PPV. While the alignment-free procedure identifies clones that are broadly consistent with the junction-based distance method, it also identifies clones with characteristics (multiple V or J gene assignments or junction lengths) that are not detectable with the *junction based distance* method.

## 1 Introduction

A defining property of the adaptive immune system is its capability to adapt to new pathogens. This property stems from the ability of B-cells to generate a broad range of antibody or Ig receptors and then to modify these receptors upon a pathogenic challenge. Each B-cell receptor (BCR) is composed of two protein chains, a heavy chain (IgH), and a light chain (IgL). The IgH is created through a somatic recombination process involving a rearrangement of three genes, termed V, D and J, coupled with stochastic insertions and deletions at the gene boundaries (i.e., between the V and D, and the D and J genes). The IgL is generated through a similar process, but without a D gene. This process provides the B-cells an initial diversity of ~ 10^7^ [3]. After a B-cell is activated through binding to an antigen and receiving appropriate secondary signals, it can undergo further diversification through somatic hypermutation (SHM). B-cells with higher affinity to the antigen are preferentially selected to further expand. This process, known as affinity maturation results in a group of clonally related B-cells. Identification of these clonally expanded groups has a tremendous biological implication, and identifying these clonal relationships are usually a first step in the analysis of B-cell repertoires. Examples for such biological studies include lineage reconstruction [4], diversity analysis [5], identification of antigen-specific sequences [6], and more [7, 8].

Our ability to analyze large B-cell repertoires had improved due to technological advances of Adaptive Immune Receptor Repertoire Sequencing (AIRR-Seq) experiments, which now allow generation of up to hundreds of millions of BCR sequences per sample. Recent studies, e.g., [9, 10, 11, 12], use AIRR-Seq to detect properties of the immune system which differentiate between healthy individuals and individuals with cancer, autoimmunity, allergies, or other diseases. Identifying these clones is an ongoing computational challenge; most current clonal identification methods rely on the strong assumption that clones should share a particular set of identifiable properties (such as V and J gene assignments, and fixed junction length).

Several methods [13, 14, 15, 16, 1, 2, 17, 18] have been proposed to address the challenge of automatic identification of clones from a set of IgH sequences. The first step is alignment and V, J gene identification of each BCR sequence using a reference of known germline V, D and J gene sequences. This step is commonly performed using IMGT/HighV-QUEST or IgBlast [19, 20]. Next, sequences are separated into different groups based on shared V and J gene assignments along with identical junction lengths, where the junction is defined as the CDR3 plus the two flanking codons. A distance (indicating a level of similarity) is computed between junctions in these smaller groups, and some form of clustering is then used to identify the clones. Various distance metrics have been used to compare the junctions, including a Hamming distance [1, 2], the Levenshtein distance [13] and metric which incorporate SHM hot- and cold-spot motifs [21, 18]. The Hamming distance is computationally efficient, but restricted to fixed sequence length comparison. The Levenshtein distance removes this restriction, but with a high computational cost and therefor does not scale to huge repertoires. Furthermore, as reported in [13] the Levensthein distance is sensitive to insertion and deletions and obtains PPV values < 96% (even when incorporating the gene assignments).

To distinguish between clonally related and unrelated sequences, earlier studies set a fixed threshold on the distance between junctions [22, 23, 24]. The authors in [14], have noticed that the distribution of distances between sequences and their nearest neighbors (distance-to-nearest) tends to be bi-modal, with a first mode corresponding to clonally related sequences and second mode corresponding to sequences without clonal relationship (singletons). Using this bi-modality, [14] proposes to set a threshold that separates the two modes. Following this observation, [1, 17] use the bi-modality of this distribution to suggest an automatic way to set the threshold. A recent method by [2] uses spectral clustering with an adaptive threshold to identify the groups of clonally related sequences.

Methods such as [1, 17, 2] identify clonally related sequences with high confidence [25]; however, their success relies on two assumptions, namely that all clone members should share the V and J gene assignments and a common junction length. The later assumption effectively ignores the possibility of SHM to introduce insertions and deletions (indels). The former premise relies on the success of a pre-processing method that aligns the sequences and assigns the V and J genes. Although most alignment methods use similar germline gene databases (such as IMGT), even for a low mutation rate of 2.5%, the assignment errors of the V and J genes are 3% [26]. Subsequently, these two types of errors will lead to a non-negligible amount of unidentified clone members (sequences), or to incorrect clonal assignments.

In this study, we present an *alignment free* approach for clonal identification; this enables us to bypass the V(D)J gene assignment step and remove the fixed junction length restriction. The *alignment free* method is based on techniques from natural language processing (NLP), specifically, we use k-mer representations and re-weight them with a term frequency inverse document frequency (*tf-idf*). The *tf-idf* is a statistical measure widely used in NLP. Next, by applying a cosine distance to the re-weighted representation, we define a *tf-idf* distance, which allows us to identify sequences derived from clonally related B-cells. This procedure does not require sequence alignment and is not restricted to sequences with the same junction length. In Fig. 1 we illustrate how the *tf-idf* distance bypasses 3 building blocks from the standard clonal assignment procedure. In the Methods section, we describe the proposed alignment free approach, and detail alternative approaches for clonal assignments. In the Results section, we evaluate the capabilities of the alignment free methods using artificial and real repertoires.

**Figure 1:**
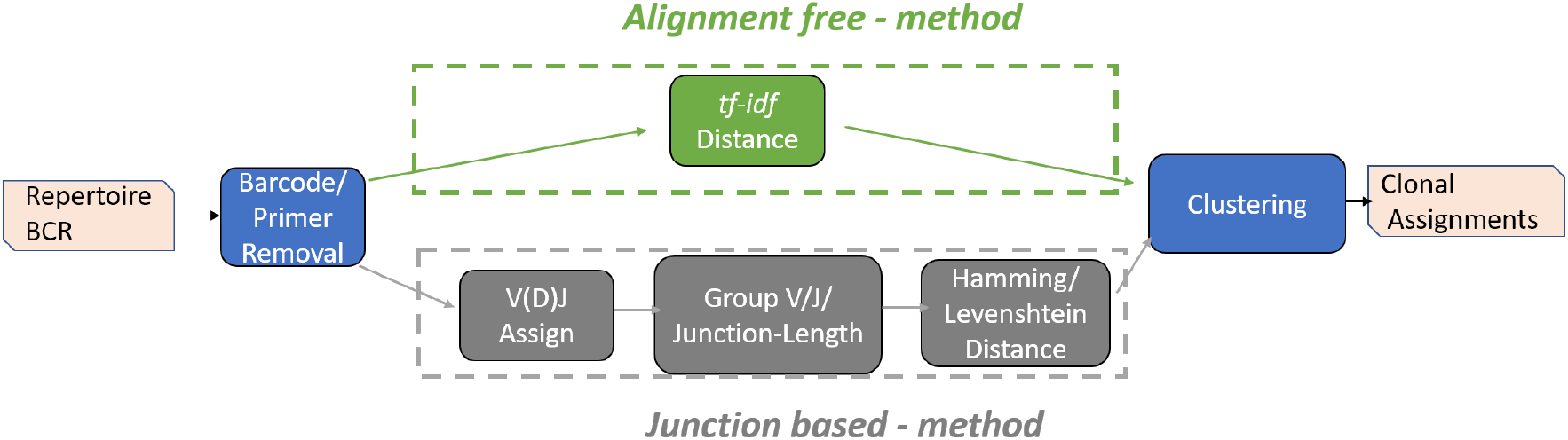
A flow diagram depicting the major steps for identifying clonally-related B-cell receptor sequences (bottom row). Given a set of BCR sequences (the repertoire), first, the primers and barcodes are removed, then V(D)J genes are assigned based on an alignment of the sequences to a database of germline genes. Sequences are grouped based on V or J gene assignments and junction length. A hamming distance is calculated on the junction regions of pairs of sequences in each of the groups separately. Finally, distances are fed into a clustering algorithm (Hierarchical [1] or Spectral [2]). Here, we propose to use a *tf-idf* based distance that bypasses the three steps prior to clustering, and is not restricted to sequences with the same V or J gene or junction length.

## 2 Methods

In this section, we describe the proposed *alignment free* approach, as well as an alternative method providing a baseline.

### 2.1 Junction Based Clonal Identification

As a baseline, we compare the performance of the proposed method to the fixed threshold, clustering-based approach proposed by [1], which has been shown to identify clonal relationships with high confidence. This approach first separates the BCR sequences into different groups that share the same V and J gene annotations, as well as a common junction length. Then, the Hamming distance is computed between all pairs of sequences from the same group, and the distribution of nearest neighbors for each sequence (also termed distance-to-nearest distribution) is analyzed to find a fixed distance threshold for single-linkage hierarchical clustering. This distance-to-nearest distribution is often bi-modal (for example, see Fig. 3), where the first mode is assumed to correspond to distances between members of the same clone and the second mode to distances between sequences from different clones. The method identifies clonally-related sequences by aggregating sequences that share a nearest neighbor distance smaller than the value (threshold) that separates the two modes of the distribution [1].

**Figure 2:**
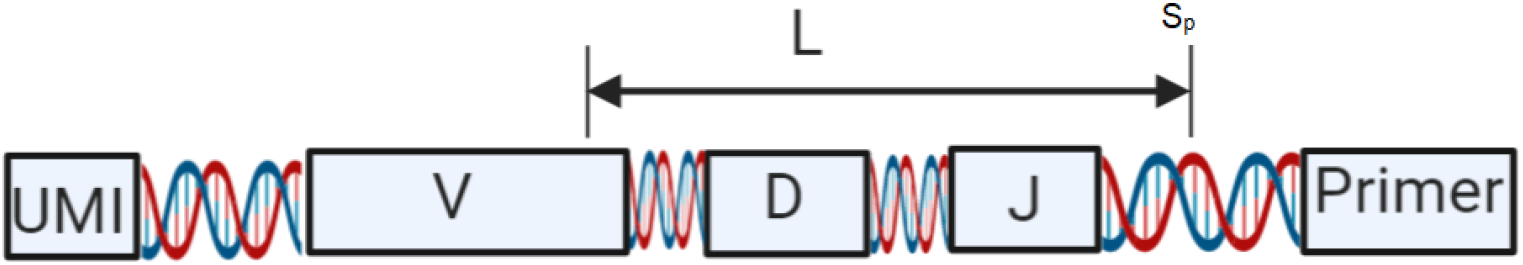
Schematic representation of a BCR sequence. The *alignment free* method uses a fixed number of nucleotides (*L*) from the 3’ end of the sequence after barcode and primer removal. Different repertoires use different library preparation procedures; thus, the starting position (*S*_*p*_) may vary and the sequence length *L* should be adjusted for each repertoire to capture the junction region.

**Figure 3:**
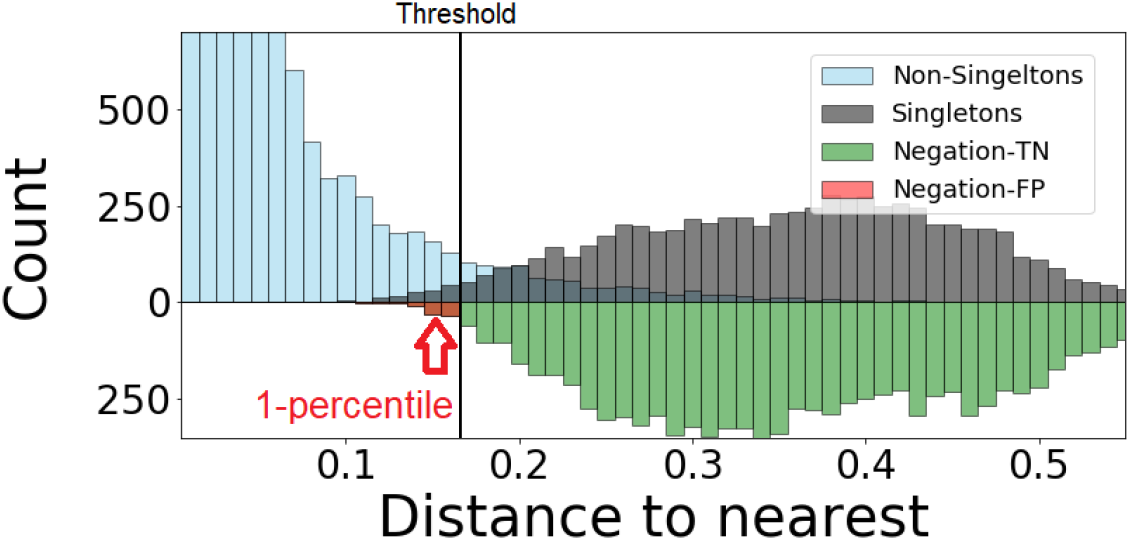
An example of a distance-to-nearest distribution based on an artificially generated repertoire, so that all clonal relationships are known. The bi-modality of the distribution is evident. Blue bars correspond to sequences that belong to a clone (non singletons), while red bars represent sequences with no clonal relatives in the data set (singletons). Green bars represent the distribution of distances to closest sequences pooled from alternative individuals (negation sequences). The distance threshold (vertical solid line)is set as the lowest 1 percentile of the negation distribution. This values aims for 1% of false positives (FP) and 99% of true negatives (TN).

The threshold is determined using a two-step process. First, the distribution of the distance-to-nearest is computed using a kernel density estimator. Next, the threshold is estimated as the first local minimum by calculating the first and second derivatives of the density estimate. As shown in [17], this is computationally expensive and empirically scales exponentially with the number of sequences. To improve run time, [17] fit a Gamma and a Gaussian distribution to the bi-modal distribution and use Maximum Likelihood to determine the threshold. This method seems to scale linearly with the number of sequences. Besides the improvement in run time, both approaches fail if the distribution is not bi-modal. This occurs if there are few or no singletons in the data, i.e., most sequences are members of clones of a size larger than one. A method to overcome this caveat was presented in [2]; the authors use spectral clustering, which allows them to identify the clones without restricting to a single threshold per repertoire.

Throughout the experiments presented in the Results section, in order to use the junction based distance method, we first apply IgBLAST with the IMGT gene references to infer gene segments and the junction location. Then, we use the Change-O-DefineClone function from SHazaM R package (version 0.1.11) [27] to define the clones.

### 2.2 k-mer Representation of Terms

A k-mer representation maps a sequence of length *L* to the set of all possible sub-sequences of length *k*. k-mers have been widely used in the context of DNA analysis due to their ability to reduce the computational effort of comparing DNA strings. The k-mer representation of a sequence is a vector with 4^*k*^ entries, where each entry corresponds to the number of times a specific k-mer is detected along the sequence using a sliding window scheme. This construction ignores the locations of particular k-mers within each sequence.

### 2.3 *tf-idf* Representation

A challenging task in natural language processing (NLP) involves comparing documents with a large and varying number of terms. One popular approach involves using a term frequency inverse document frequency (*tf-idf*) [28]. The *tf-idf* weighting scheme aims to emphasize the rare and hopefully meaningful terms and reduce the influence of common terms. The *tf* counts the number of term appearances in the document, while *idf* measures the importance of the term by counting its appearances in the corpus. A number variants of the *tf-idf* have been proposed in the literature, for examples see [29, 30].

In this study we adapt the *tf-idf* to reweigh k-mer based representation of BCR sequences. The term-frequency *tf*_*s*_(*k*) is a count table of the amount of k-mers *k* ∈ *K* present in each sequence *s* ∈ *S*. The *tf* is reweighed using the inverse document frequency which is defined as 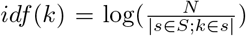, where *N* is the total number of sequences and the denominator is the total occurrences of specific k-mer *k* across all the *S* sequences. The *tf-idf* is then defined as *tf-idf*_*s*_(*k*) = *tf*_*s*_(*k*) · *idf* (*k*).

### 2.4 Fast Cosine Distance

To compare the *tf-idf* representations of sequences *s* and *s*′ we choose the widely used cosine distance. The cosine distance is defined as one minus the normalized inner product between two *tf-idf* vectors, given sequences *s* and *s*′ it is computed by

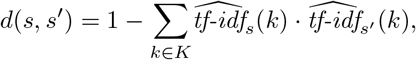

where 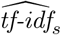 is the *L*_2_ normalized *tf-idf* representation of sequence *s*. In practice the *tf-idf* representation is sparse and clone assignment only requires identification of the most related sequences in terms of proximity (few hundreds, or thousands depending on the size of the clones in the dataset). We can exploit these two properties using a python implementation^1^ that reduces the computational cost involved in evaluating the *tf-idf* cosine distance.

### 2.5 Automatic Clonal Distance Threshold Determination by Negation

As in [1, 17] we use the properties of the distance to nearest distribution to identify the threshold for cutting the hierarchy of distances. Instead of using the bi-modality, which has a high computational cost, we take a different approach; we propose to find the threshold by negation. The idea is described as follows: given a set of sequences taken from one individual, we introduce a set of sequences randomly sampled from multiple alternative individuals (negation sequences); then, we compute the distribution of distances between negation sequences and their closest counterpart within the individual. Finally, we set the threshold such that a fraction of the distances to negation sequences that are below the threshold is *δ* ≥ 0. By definition, clones can not span multiple individuals. Therefore, by choosing a fixed value for *δ* > 0 (e.g. *δ* = 0.01) we allow a fraction of false-positive rate roughly equal to *δ*. This heuristic aims for high specificity, which is approximately 1 − *δ*. We present an example of such distribution along with the estimated threshold in Fig. 3.

### 2.6 Simulation of Clonal Expansions

To generate artificial repertoires with known clonal relationships, we first select clone representatives from B-cells collected by [24] and filtered to maintain only naive sequences from healthy individuals as in [31]. Next, we infer tree topologies of each clone by applying Change-O-buildPhylipLineage (version 0.4.5 [27]) to data from multiple individuals collected in [7]. Finally, new artificial samples are generated by randomly adding mutations based on the learned topologies using shmulateTree from the SHazaM R package (version 0.3.0 [27]). We repeat this process using repertoires from four subjects collected in multiple sclerosis (MS) study [32]. The corresponding four datasets which contain samples from lymph nodes and are denoted as MS2, MS3, MS4, and MS5 with around 100K, 150K, 200K, and 200K sequences, respectively. Using the sequences from the four individuals we generate 74 simulated datasets with ~ 30*k* sequences each. To support the diversity of the artificial datasets, we analyzed the properties of the resulting 74 datasets and observed that the distributions of sequence and junction lengths both follow Gaussian-shaped distributions with means of 521 and 58 nucleotides, respectively. Moreover, as evident in Fig. 4, the generated repertoire has a wide range of sequence and junction lengths and diverse clone sizes.

**Figure 4:**
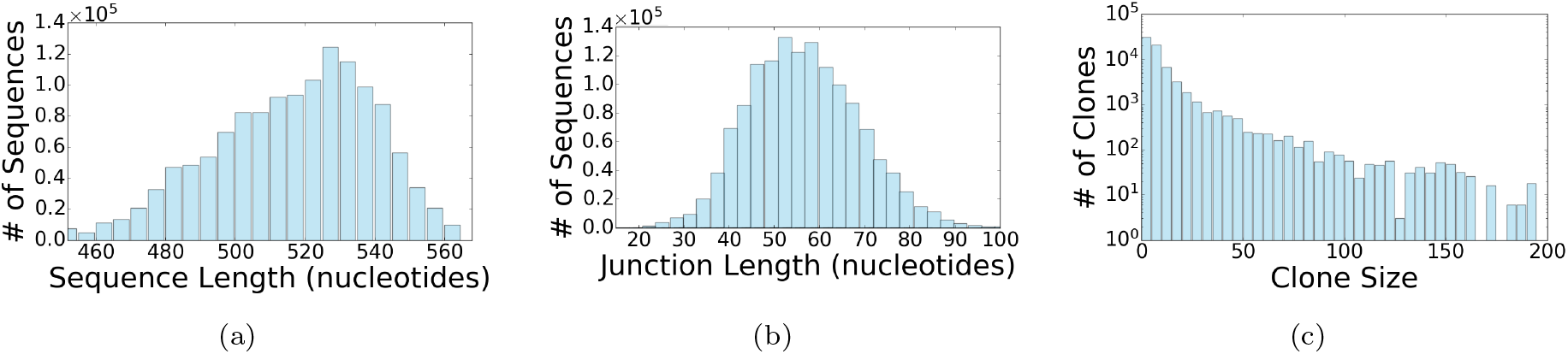
Statistical properties of the 74 artificially generated datasets. (a) The distribution of sequences length. (b) The distribution of junction lengths. (c) The distribution of clone size (number of unique sequences).

## 3 Results

We propose an *alignment free* method for identification of clonally related sequences. The approach relies on a *tf-idf* based distance that is invariant to translations of the sequence and is not restricted to the comparison of sequences with the same length. The *alignment free* method does not require alignment of sequences to germline V, D or J genes. The only pre-processing required is removing primers and truncating the sequences to a fixed length from the 3’ end. Here, we evaluate the *alignment free* approach using both simulated and real datasets. For the simulated repertoires, the correct clonal assignments are known; this allows us to compute the sensitivity, specificity, and positive predictive values (PPV). We define sensitivity as the ratio between pairs correctly identified as clonally related and the total number of pairs which truly belong to the same clone. We calculate PPV as the ratio between correctly assigned pairs and the total number of clonal assignments. For specificity, we compute the fraction of identified unrelated sequence pairs among all truly unrelated pairs. To further evaluate the alignment free using real repertoires, we focus on a set of published BCR repertoires from lymph nodes of four multiple sclerosis (MS) subjects referred as MS2-MS4.

### 3.1 *idf* normalization using a fixed sequence length improves sensitivity and specificity

We start by evaluating how the *idf* normalization influences the performance of the *alignment free* approach. The *alignment free* method uses nucleotide *k*-mers reweighed by a *tf-idf* normalization to find a translation-invariant representation for BCR sequences. We hypothesize that this representation captures sufficient information to allow efficient clonal identification. The term-frequency *tf*_s_(*k*) counts the number of occurrences of each k-mer; thus, the total count of terms per sequence *s* is affected by the sequence length. As shown in Fig. 4(a) the length of the sequences range between 460 and 560 nucleotides per sequence. In practice, the bounds of this range can vary depending on the experimental library preparation method although there will always be a distribution of lengths resulting from the V(D)J recombination process. Such variability in the sequence length, in turn, translates to a variability in the amount of *k*-mers present in each sequence. As been shown in [33], such variability may bias the *tf-idf* based representation. Specifically, applying cosine similarity favors retrieval of short documents (sequences) over long ones.

In [33], the authors propose to use a pivoted normalization to compensate for the length effect. Here we can exploit the known structure of the BCR sequences to propose an alternative solution by defining a representation that uses a common number of nucleotides for all sequences. This can be obtained by truncating each sequence using a fixed number of nucleotides (*L*) starting from the 3’ end. The length of the sequence should be sufficient to cover the J segment, the junction region, and part of the V segment. See Fig. 2, in which we illustrate this truncation. We expect that a good truncation should cover the junction region, as it is highly diverse and has been used as a signature for identifying clonally-related BCRs in several studies [1, 2].

Given a sequence ***S***^*i*^ = [*S*^*i*^(1), *S*^*i*^(2), …, *S*^*i*^(*N*^*i*^)], where *N*^*i*^ is the length of the *i*^*th*^ sequence and *S*^*i*^(*x*) indicates its *x*^*th*^ nucleotide, we define the truncated sequence of size *L* (number of nucleotides) as 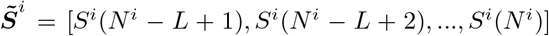. First, from the 74 artificial datasets, we use a subset of 20 repertoires and seek an optimal value of *L*, the length of the sequence used for the *tf-idf* representation. The remaining 54 datasets will be used later to evaluate the method’s performance. To compare the performance across different *L* values, for each value of *L* in the range [100, 300] we first tune the clustering distance threshold to obtain 99% specificity. Then we evaluate the sensitivity and PPV. Based on statistics of aligned sequences from the MS dataset, we expect that this range of lengths will be sufficient to cover all of the J segment along with the adjacent junction regions (see Fig. 6). As depicted in Fig. 5(a), when *L* is in the range [120, 150] performance is peaked. For the rest of the experiments, we use *L* = 150 nucleotides from each sequence.

**Figure 5:**
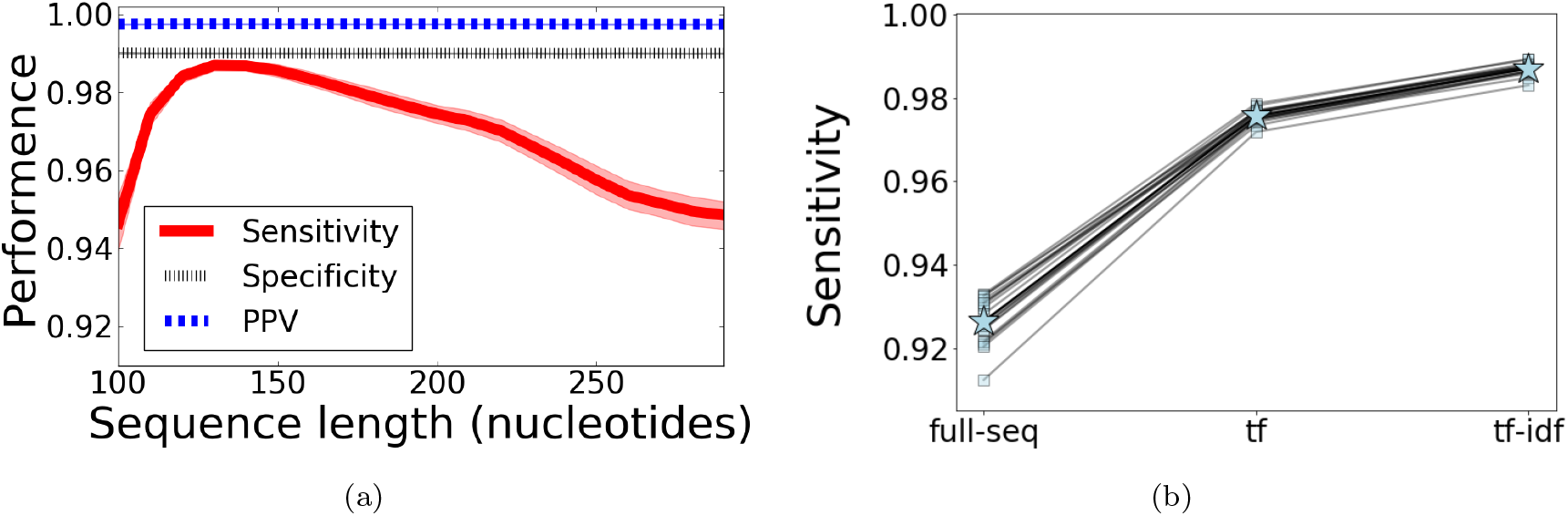
(a) Performance analysis vs. the number of nucleotides used from each sequence (*L*). The line represents mean sensitivity, specificity and PPV, while the shaded color displays the standard deviation over 20 different artificial repertoires. (b) Performance comparison for three different settings; *tf-idf* applied to the full sequence (*full-seq*), *tf* applied to *L*=150 nucleotides from each sequence (*tf*) and *tf-idf* applied to *L*=150 nucleotides from each sequence (*tf*).

**Figure 6:**
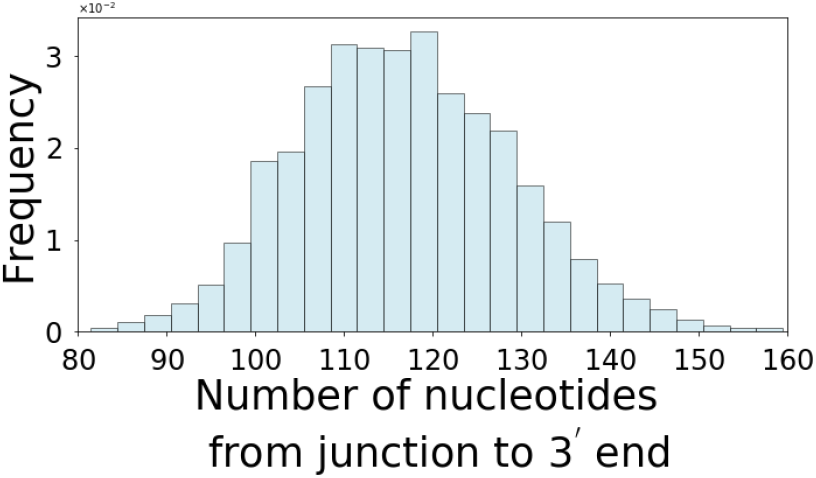
150 nucleotides is sufficient to cover the junction in most sequences. The number of nucleotides from the start of the junction to the 3’ end was calculated for each sequence of 20 artificial repertoires.

Next, we compare the performance of the *alignment free* method using three different variants of the *tf-idf* representations. The first term entitled *full-seq*, in which the *tf-idf* is applied to the full sequence (with variable length). The second, referred as *tf* in which only the *tf* normalization is applied to a fixed part of the sequence with length *L* = 150. The last, named *tf-idf*, inputs a fixed part of the sequence as *tf* does, but uses both the *tf* and *idf* normalizations. As in the field of NLP, we speculate that the *idf* normalization will help by up weighting the unique k-mers, which represent the diverse junction region derived from V(D)J recombination as well as unique SHM that are shared by clonal relatives. At the same time, the *idf* should down weight the common k-mers, which correspond to the unmutated V, D, and J gene sequences. As in the previous experiment, we use a threshold that achieves 99% specificity and evaluate the sensitivity of the *alignment free* method. By applying the *tf-idf* to *full-seq* we obtain ~ 93% sensitivity, using a fixed part of the sequence improves the performance to ~ 99%, while without the *idf* normalization the sensitivity is ~ 97.5% (see Fig. 5(b))

### 3.2 The alignment free has high sensitivity, specificity, and PPV

Next, we compare the overall performance of the *alignment free* method to the performance of a widely-used *junction-based distance* method [1] (see Supporting material for more details) using the remaining 54 simulated repertoires. Comparing to a state-of-the-art-method would allow us to evaluate whether the alignment step (along with gene assignment and sub-grouping) is necessary for accurate clonal assignment.

To compare the two methods in terms of sensitivity, specificity, and PPV, we first apply the *junction-based distance* method (describe in the Methods section) and evaluate performance. For the *alignment free* method we tune the threshold using the negation approach aiming for a specificity value of 0.99, here the negation sequences are naive BCRs from [24]. We assign clones based on the estimated threshold and evaluate the *alignment free* method’s sensitivity, specificity, and PPV values. The *alignment free* method achieves roughly 4% higher sensitivity compared with the *junction-based distance* method, while maintaining similar specificity and PPV (values presented in Fig. 7(a)).

**Figure 7:**
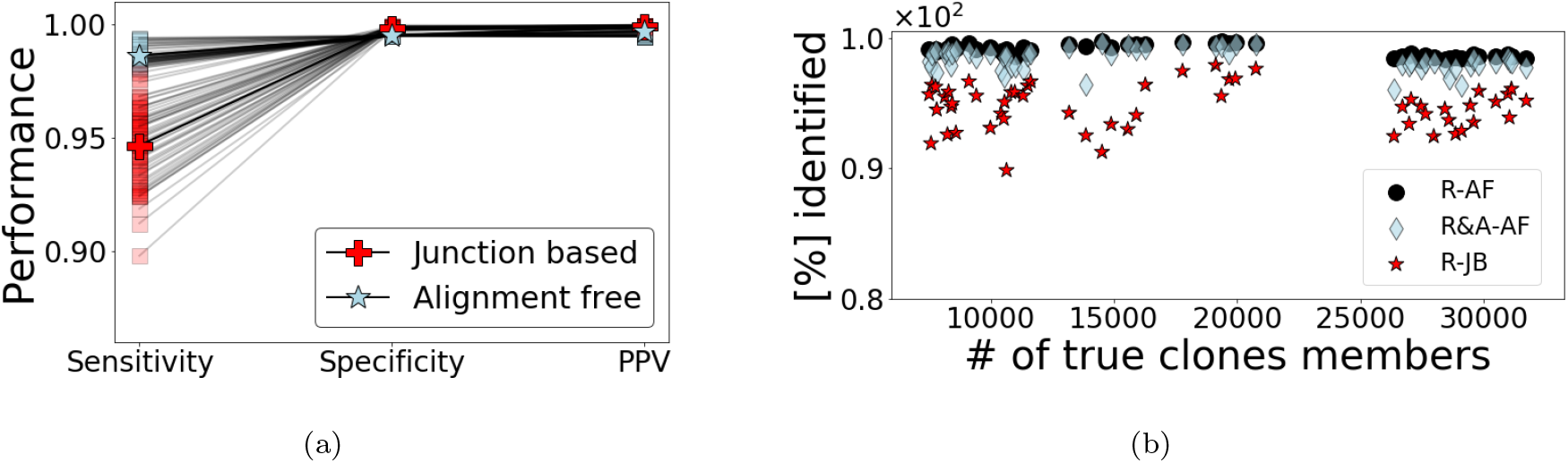
(a) Performance comparison of the *alignment free* method against the *junction-based distance* based on 54 artificial datasets. We evaluate each approach by measuring whether assignments were correct for all input sequences. (b) The percentage of sequences assigned to their correct clone (y-axis) sorted as a function of the true number of clone members in each of the 54 datasets (x-axis) by the *alignment free* method (RA-F), the same method but evaluating portion of clone members with correct V and J gene assignments (R&A-AF). Here the gene assignments are performed as a final step and only based on one clone representative. Clone representative is selected randomly from the sequences that do not contain non-ACGT characters (gaps or N’s). The red stars represent the *junction-based distance* approach with V and J gene assignments via IgBlast (R-JB).

One advantage of the *alignment free* method is that it does not require running the full repertoire though a V(D)J assignment program like IgBlast. However, once a clone is identified, it is still important to determine the V, D and J assignments for biological interpretation. To achieve this, we randomly select a clone representative that does not contain non-ACGT characters (gaps or N’s) and run it trough IgBlast for gene assignment. Following this procedure, gene assignment is performed as a final step and is only applied to a subset of sequences.

To evaluate whether this procedure is beneficial compared with the *junction-based distance* method (see Fig. 1), we compute the amount of clone members recovered with correct V and J assignments based on both pipelines. For each dataset, we compare the total number of members in expanded clones (True), the number of expanded clone members recovered by the *junction-based distance* method [1] (R-JB), the number of expanded clone members retrieved by the *alignment free* method (R-AF) and the number of expanded clone members with correct V and J assignments by the *alignment free* method (R&A-AF).

Identification of the clonal membership before performing V(D)J assignment increases the portion of identified (true) clone members by more than 4% (see Fig. 7(b)). This gap could be explained by the fact that the *alignment free* method can still recover clones with incorrect alignments or with non-ACGT characters (gaps or N’s). This experiment demonstrates that not only the alignment step could be avoided, but also that more sequences are retrieved if the alignment is performed as a final step based on one clone representative.

[1, 17] observed that the ability to correctly identify clonal relationships drops for BCRs with shorter junction lengths. This drawback was improved in [2] using spectral clustering with an adaptive distance threshold. Here, since we use a single (fixed) threshold to identify clones, we expect that the performance of the *alignment free* method will deteriorate for sequences with shorter junctions. To evaluate how the length of the junction affects the *alignment free* method, we apply it to the simulated repertoires, and compute the sensitivity, specificity and PPV for three different ranges of junction length (*L_J_*), namely (0, 30), [30, 60] and (60, 90]. These ranges were selected as they separate the data to approximately equal size subsets. As evident in Fig. 8, the *alignment free* method indeed achieves higher performance when focusing on sequences with longer junctions.

**Figure 8:**
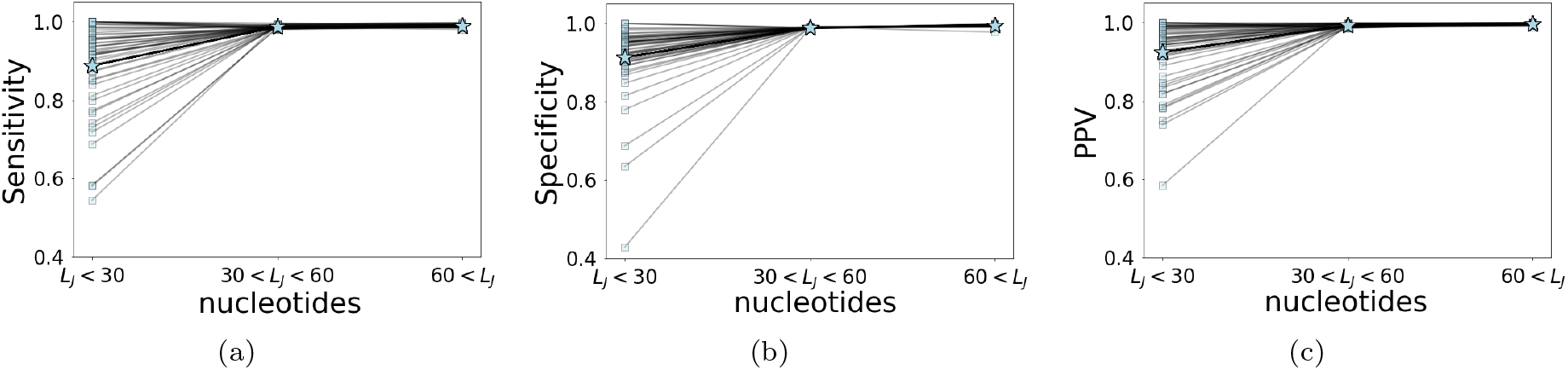
The *alignment free* approach performs better for sequences with longer junctions. (a) Sensitivity, (b) specificity, and (c) PPV as a function of the junction length (*L*_*J*_) evaluation using 54 artificial repertoires. Square marker correspond to performance on each individual dataset, while star markers represent the mean value.

### 3.3 The alignment free method has low false-positive rates on real repertoires

In this section, we compare the *alignment free* approach to the *junction-based distance* method on a set of experimentally-derived human repertoires. We use previously published sequencing data from four MS subjects [32] (MS2, MS3, MS4, and MS5). In three of these datasets (MS3-MS5), the automatic distance threshold identification applied by the *junction-based distance* method (SHazaM-findThreshold [27]) fails, as the distance-to-nearest distribution of these samples is not bimodal. Therefore, following [1] for the *junction-based distance* method, we use the threshold that was estimated based on MS2 to identify clones in MS3-MS5. The same lack of bimodality appears when evaluating the distance-to-nearest of the *tf-idf* representation. To estimate the threshold, we use the negation method (explained in the Methods section) with naive sequences from [24]. The negation threshold is tuned to obtain ~ 0.99 specificity based on MS2. Finally, to have the *alignment free* method consistent with the setting in [1], we use the same threshold from MS2 for clonal assignments in MS3-MS5.

To compare the clonal assignments of the *alignment free* to the assignments made by the *junction based* method, we compute the normalized mutual information (NMI) between the clonal assignment of both methods. The NMI measures how well the two clonal assignments predict one another, it has a highest possible value of 1. As appears in Table 2, the NMI between assignments for all the individuals is high (> 0.93), which indicates a high consistency between both clonal assignments methods. As a complementary evaluation, we show that the distribution of clones size is consistent across all four individuals (see Fig. 9).

**Figure 9:**
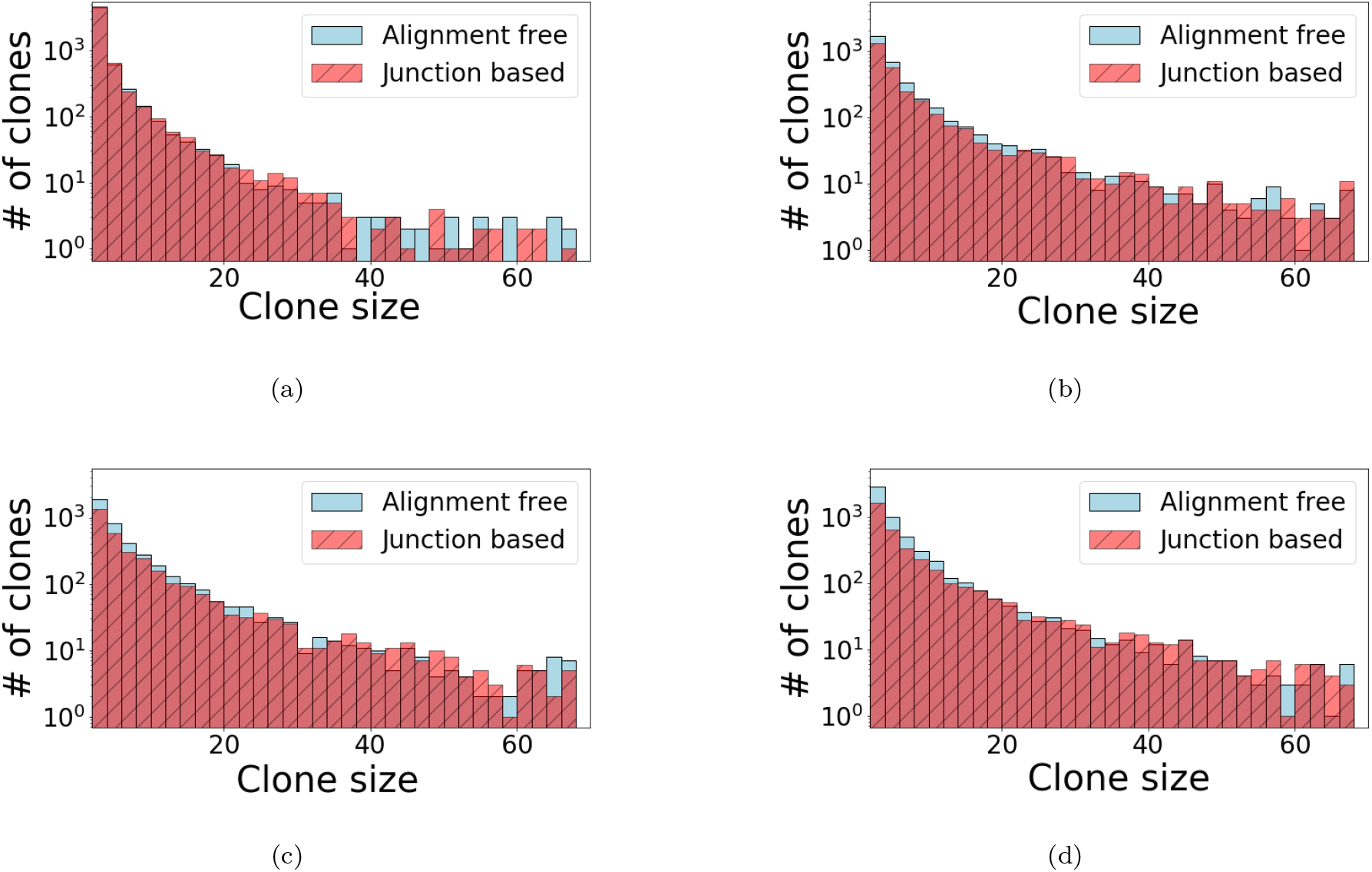
High clustering consistency between the *junction-based distance* [1] and *alignment free* methods using sequences from four individuals with MS [7]. A comparison between the distributions of clone sizes identified by both methods, (a)-(d) correspond to MS2-MS5.

To evaluate the false-positive rates of the *alignment free* method on real data, we rely on the fact that clones cannot be shared across different individuals. This is because clonally related cells develop from a common ancestor with a single V(D)J germline rearrangement. The false-positive rates are evaluated by first computing the *tf-idf* base distance-to-nearest sequences across individuals (see Fig. 10. Next, we applied the negation based automated threshold estimation (described in the Methods section) using data from [24]. Here the threshold is tuned based on the 1 percentile distance to negation sequences (i.e. aiming for 99% specificity). Clones are defined by cutting the hierarchy based on the estimated threshold, and we count the fraction of clone members form mixed individuals. This portion provides an estimate of the false-positive rates in the MS data. We apply this procedure to all pairs of individuals in the MS data and observe that the false-positive rates are lower than 0.5% (see table 1).

**Table 1:** The *alignment free* approach was applied to pairs of individuals (MS2-MS4 from [7], rows and columns) and the percentage of sequences predicted to be part of clones that span individuals was calculated. These clonal relationships are considered false positives, as clones cannot be shared across individuals.

|  | MS-2 | MS-3 | MS-4 | MS-5 |
| --- | --- | --- | --- | --- |
| MS-2 | NA | 0.34 [%] | 0.23 [%] | 0.27 [%] |
| MS-3 | 0.34 [%] | NA | 0.032 [%] | 0.012 [%] |
| MS-4 | 0.23 [%] | 0.032 [%] | NA | 0.014 [%] |
| MS-5 | 0.27 [%] | 0.012 [%] | 0.014 [%] | NA |

**Figure 10:**
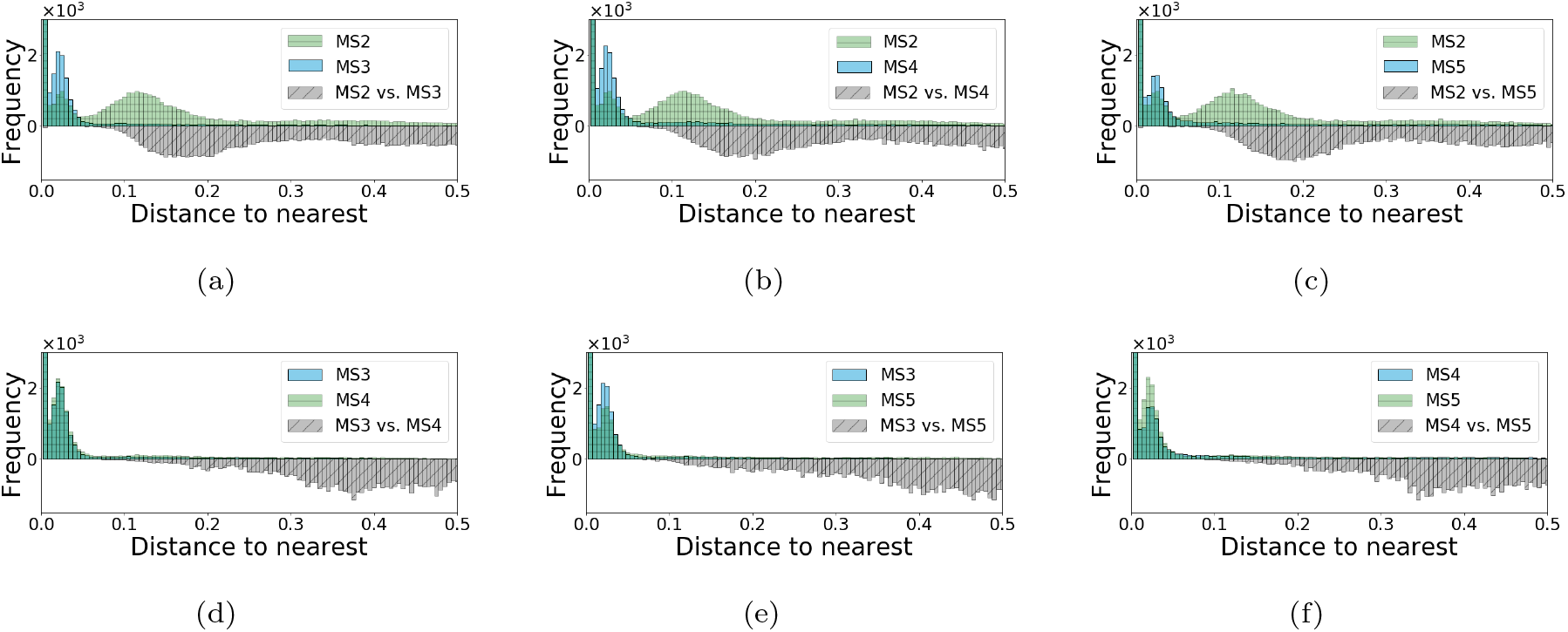
Distribution of distance-to-nearest in the *tf-idf* space. Distances were computed between sequences within one subject (above the *x*-axis) and between pairs of subjects (below the *x*-axis), for all pairs of subjects in the MS dataset.

### 3.4 The alignment free method identifies novel clonal relationships

As the *alignment free* method is not restricted to a fixed junction or common V or J gene assignments it has the potential to retrieve novel clonal relationships. Here, we evaluate whether the *alignment free* method identifies such novel relationships in real data.. Specifically, we identify clones with multiple V and J gene assignments (when such assignments are made on a sequence-by-sequence basis) or non-unique junction lengths. First we turn our attention toward estimating positive predictive value (PPV) of the *alignment free* when used on real repertoires.

Clone members evolve from the same germline; therefor, they should all share the same V and J genes. We use this property to bound the PPV value of the *alignment free* method. Based on repertoires from the MS study, we observe that the number of clones identified with non-unique J genes is < 0.06% of the total number of identified clones (see the full comparison in table 2). More significantly, we found that even though we do not use the junction length and only use a small portion of the V gene, the percentage of sequences with non-unique V genes or junction lengths is also low (< 0.12%). If we consider these clonal relationships as errors, they provide an upper bound for the PPV in the MS dataset. However, it is possible that these clonal assignments are correct. For example, sequences with different V or J gene annotations in the same clone could result from incorrect assignments by IgBlast. Such relationships are possible, as the accumulation of SHM can make a BCR derived from single V or J genes seem to stem from distinct V or J genes. Clonal relatives may also have different junction lengths due to the occurrence of indels, which can accumulate as a part of normal SHM [34].

**Table 2:** Properties of clones identified by the alignment free method. NMI refers to the Normalized Mutual Information between clonal assignments made by the alignment free method compared to the *junction-based distance* method. We also determined the percentage of clones with members expression non-unique V or J genes, or non-unique junction lengths.

|  | MS-2 | MS-3 | MS-4 | MS-5 |
| --- | --- | --- | --- | --- |
| **NMI (with JB)** | 0.98 | 0.96 | 0.96 | 0.93 |
| **Non unique V [%]** | 0.008 | 0.008 | 0.006 | 0.002 |
| **Non unique J [%]** | 0.038 | 0.036 | 0.054 | 0.014 |
| **Non unique Jun. [%]** | 0.11 | 0.053 | 0.027 | 0.025 |

To further evaluate whether we have identified true clonal relatives whose initial V or J gene assignments were incorrect or if the non-unique V or J gene assignments are, in fact, false positives, we use a normalized Levenshtein distance [35]. The Levenshtein distance (also termed edit distance) finds the minimal number of single edits required to change one sequence to the other. In contrast to the *alignment free* approach, this comparison takes into account the exact locations in the BCR, and it therefore more accurate than the *tf-idf* based distance. We note that a direct application of Levenshtein distance to all pairs of sequences is computationally exhaustive. Specifically, for a repertoire of size *N*, with sequences of length *m* complexity is *O*(*Nm*)^2^.

In Fig. 11, we first focus on clones with non-unique V or J gene assignments, we present a histogram of a normalized Levenshtein distance (see definition in the Supporting material) between the pairs of closest sequences with different V or J gene assignments. As a background test, we select random groups of sequences, consisting of the same number of sequences as in the retrieved clones. All sequences in such a background group share a V and J gene (majority group) except one sequence with a different gene (minority sequence). Then, we compute the smallest distance between the minority sequence and all sequences in the majority group. The background distribution represents a histogram of such nearest distances (see Supporting material for more details on this procedure). As evident from Figs. 11(a) and 11(b) the majority of the clones composed of sequences with non-unique V or J gene assignments identified by the *alignment free* method have a low distance-to-nearest (relative to the background distribution) when comparing to randomly chosen sequences with different V or J assignments. This supports that a non-negligible portion of the identified sequences are indeed clonally-related, and may have diverged due to SHM. Furthermore, in the Supporting material, we recompute the distance-to-nearest between the junction part of the sequences, and observe similar low values (relative to the background distribution).

**Figure 11:**
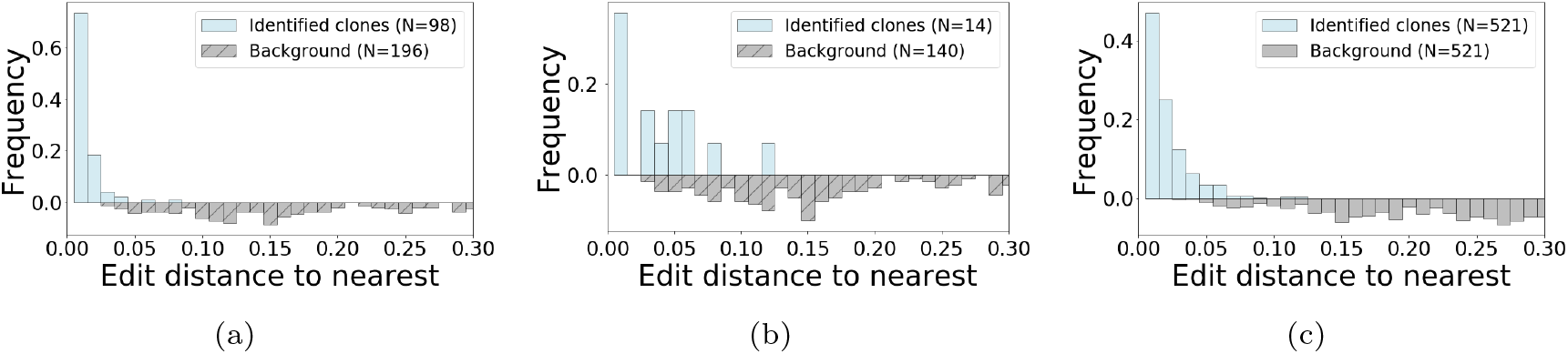
Evaluation of clones with non unique V gene, J gene or junction length. We use the normalized Levenshtein distance (see definition in the Supporting material) to asses clones with non-unique V, J, or Junction length. (a) Distribution of Levenshtein distance from each sequence to its nearest non-identical neighbor within clones with non-unique J gene assignments (b) or non-unique V gene assignments, or (c) non unique junction lengths.

Next, in Figs. 11(c) we show that the distribution of distance-to-nearest among clones with multiple junction lengths is also low (compared to the background distribution), which explains why these sequences were pulled together. A complementary junction based comparison shows that some of these groups might contain members with highly diverse junctions (see Supporting material). To further evaluate clones with non unique junction lengths, we study the structure of their phylogenetic trees. Lineage trees were constructed for each clone using Change-O-buildPhylipLineage (version 0.4.5 [27]. The inferred tree topologies (see Fig. 12) show that these clones have a non-negligible amount of shared mutations relative to the germline sequence. This is a positive indication that even though these clones have sequences with different junction lengths, they are likely to have evolved from a common mutated ancestor. Furthermore, the average number of minimal shared mutations across the clones with multiple junction length is 7.5 (full distribution appears in Fig. 12(d)). These results demonstrate the potential of the *alignment free* method in retrieving clonal relationships between sequences with different junction lengths, or sequences that were assigned to different V or J genes when analyzed individually.

**Figure 12:**
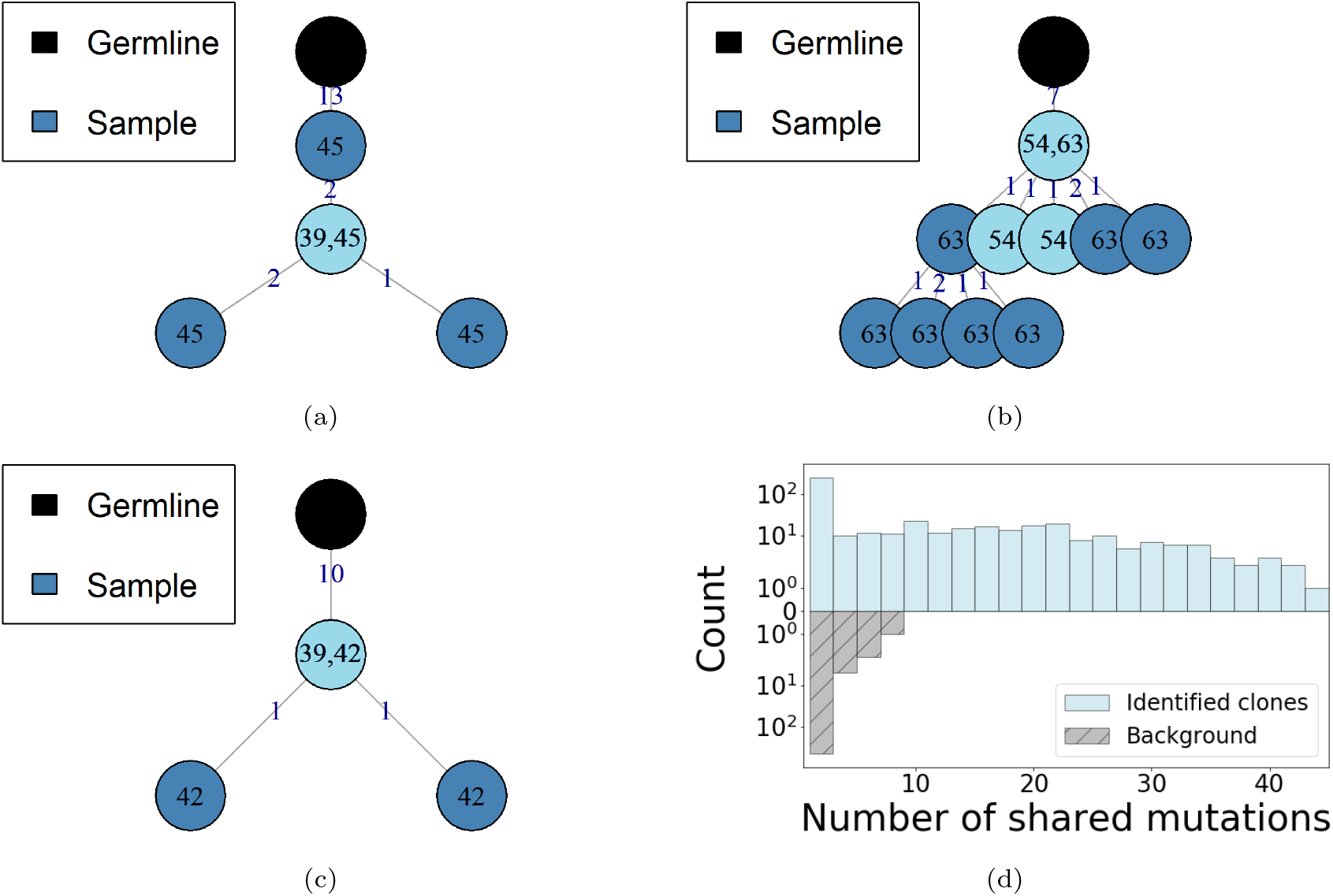
Tree topologies inferred from clones that include members with multiple junction lengths as determined by the *alignment free* method (a), (b) and (c). Branch numbering indicates number of shared mutations, node numbering represents the length of the junction. The germline sequence is colored in black and light blue represents the minority junction length sequences. (d) The distribution of the minimum number shared mutations between sequences with different junction lengths within each clone. The background distribution is computed by sampling random groups of sequences that share the same V and J genes but different junction lengths.

## 4 Conclusion

B-cells play a crucial role in the active immune system. Their ability to recognize and efficiently respond to antigens relies on two diversification mechanisms. The first occurs at an early stage of maturation and acts by joining V and J genes (and D gene for heavy chains) to create a functional antibody receptor. The second part of diversification occurs in the germinal center trough SHM; this step generates a diversified group of clonally related B-cells. A critical step in the analysis of high throughput B-cell receptor sequencing data is the identification of groups of such clonally related B-cells.

We have presented an *alignment free* method for clonal identification. The approach uses a nucleotide k-mer representation to define a term frequency-inverse document frequency (*tf-idf*) based distance. This distance is invariant to the exact locations of the k-mers in the BCR sequence; thus, it allows us to bypass the V(D)J alignment step. A second advantage of the *alignment free* method is that we can identify clonally-related sequences with multiple junction lengths, which can be generated though the accumulation of indels and can be important in affinity maturation to some pathogens. To evaluate the capabilities of this new procedure, we generate simulated repertoires with known clonal relationships between all of the sequences. Using these repertoires, we demonstrate that the *alignment free* method has high sensitivity, specificity, and PPV. Furthermore, our results suggest that by performing the V(D)J gene assignment after clonal identification, more clone members are retrieved.

We apply the *alignment free* method to real repertoires collected from four MS subjects. These repertoires lack a correct known clonal assignment; nonetheless, two observed properties suggest a low false positive rate; a low frequency of identified clones containing sequences with different V or J gene assignments, and a low incidence of clones shared across individuals. These features are expected to be enriched among potential false positive relationships. However, it is likely that at least some of them result from incorrect V or J gene assignments by IgBlast, in which cases the alignment free method identifies clone members which would be lost by other approaches.

The library preparation of the MS data is fairly typical; in other experimentally-generated repertoires the precise location of primers may differ and the number of nucleotides required for the *tf-idf* representation (*L*) may vary. For more information, see Fig. 2, in which we describe the different parts of the BCR and discuss the variability that may arise from different library preparation processes. Our experiments demonstrate that using *tf-idf* and restricting to a fixed part of the sequence improves sensitivity, specificity, and PPV values. Nonetheless, the fixed-length used (*L*) should be adapted to the library preparation technology of each repertoire. Specifically, *L* should be large enough such that it covers the full junction region (counting nucleotides from the 3’ end). In Fig. 6 we present the distribution of the number of nucleotides required to cover the junction region based on 20 artificial repertoires.

The final step of the *alignment free* method requires clustering sequences into clonal groups. Here, we identify these groups by thresholding the dendrogram of distances between sequences. We implement this hierarchical clustering procedure using a fixed threshold, and we optimize this threshold using negation sequences. By computing the distance-to-nearest negation sequences, we optimize a single threshold to obtain high specificity. As demonstrated using simulated repertoires, the performance of the *alignment free* using a single threshold deteriorates when focusing on sequences with short junctions. A natural extension of this work, could alleviate this shortcoming by considering multi-scale thresholds. One example for such solution was presented in [2], where the authors use spectral clustering with an adaptive threshold to identify the clones.

Overall, we have developed an *alignment free* clonal identification method using tools form natural language processing. We demonstrate using artificial and real repertoires that the *alignment free* compares to state-of-the-art distance-based methods, in terms of sensitivity, specificity and PPV. This shows that the fundamental task of identifying clonal groups does not have to rely on V or J gene assignments. Finally, as the method is capable of identifying clonally-related BCRs with different junction lengths it represents an important improvement in clonal assignments for AIRR-seq analysis.

## Appendix

### 6 Normalized Levenshtein Distance

The Levenshtein distance is a distance metric between two DNA or protein sequences. It measures the minimum number of single character (base pairs) edits require to change one sequence to another. Each edit is either an insertion, deletion or substitution. Computing the Levenshtein distance, requires finding the optimal alignment that requires the minimal number of edits, and therefore is considerably more expensive computationally. In Fig. S1, we illustrate the Levenshtein distance computation.

**Figure S1:**
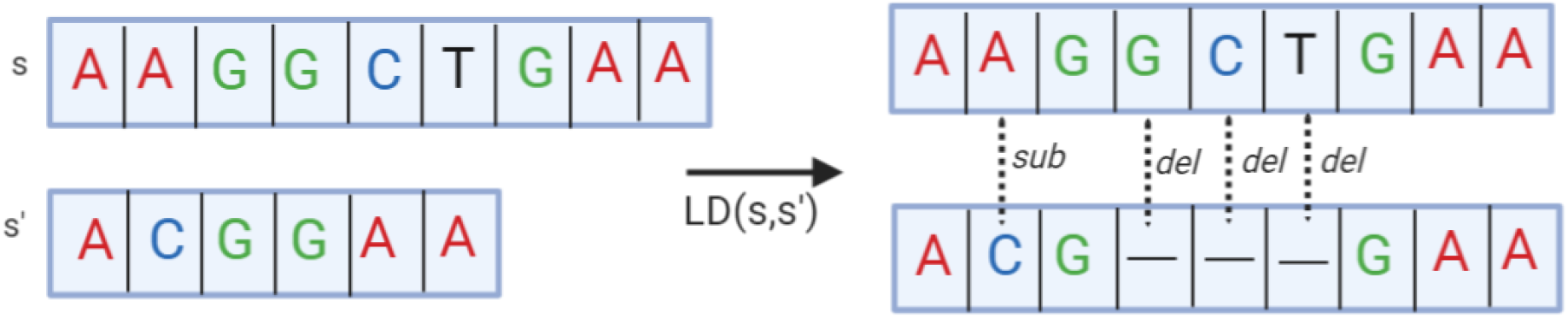
The Levensthein distance between sequence *s* and *s*′ is determined by the minimum edits required to transition form one sequence to the other. In this example three deletion (d) and one substitution (s) are the minimal number of edits required, thus the Levenshtein distance is 4.

The Levenshtein distance allows us to compare two sequences with different length. To reduce bias caused by length differences, [36] propose a normalized Levenshtein distance that incorporates the length of both sequences. The normalized Levenshtein distance is defined as

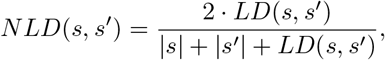

where *LD*(*s, s*′) denotes the Levenshtein distance and |*s*|, |*s*′| are the lengths of sequence *s* and *s*′ respectively.

### 7 A deeper look into clones with multiple V, J or junction length

In the main text, we have analyzed clones identified using the *alignment-free* method with non unique V, J gene assignments or with multiple junction lengths. We have computed the histogram of the distance-to-nearest between sequences with such non unique properties. A schematic description of the distance-to-nearest evaluation of such clones appears in Fig. S2. The histograms presented in Fig. 11 (main text) are based on the Levenshtein distance metric applied to pairs of full sequences. This includes the V, J segments and the junction. Next, we repeat the procedure presented in Fig. S2, but apply the Levenshtein distance only to the junction part of the sequence. In Fig. S3, we present the distributions of distance-to-nearest within clones with non unique characteristics and background groups. The distributions of clones with non unique V and J (3(a) and 3(b)) provide yet another support that the majority of these obtained clones are enriched with true clonal expansions.

Finally, in Fig. S4, we present a multi-sequence alignment of all the sequences in the clones which are presented in Fig. 12 (main text). The shared mutations in these alignment demonstrate that sequences with multiple junction length can belong to the same clone and result from clonal expansion.

**Figure S2:**
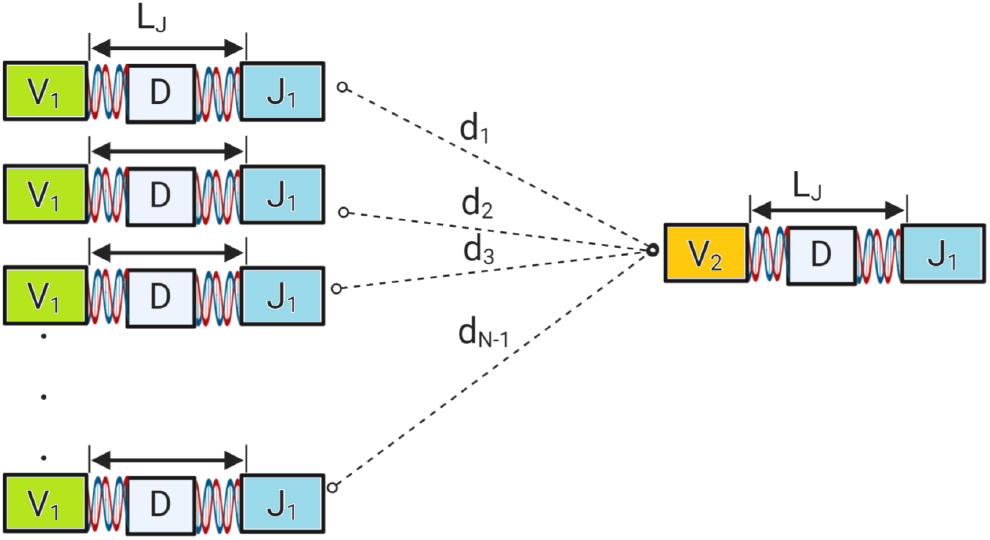
Here we illustrate the distance-to-nearest computation used in Fig. 11 (main text). The group of *N* − 1 sequences (majority group) share the same V,J genes assignments and have the same junction length *L*_*J*_. In this example, one of the sequences has a different V assignment (minority group). We compute all distances between the minority and majority groups *d*_1_, …, *d*_*N* − 1_ and compute the histogram of the minimal Levenshtein distances min(*d*_1_, …, *d*_*N* − 1_) over all clones. As a background distribution, we generate artificial groups of sequences that share the same V, J and junction characteristics, and include an additional sequence that differs by one of these characteristics. The number of sequences in each such artificial group is set using the actual sizes of the clones.

**Figure S3:**
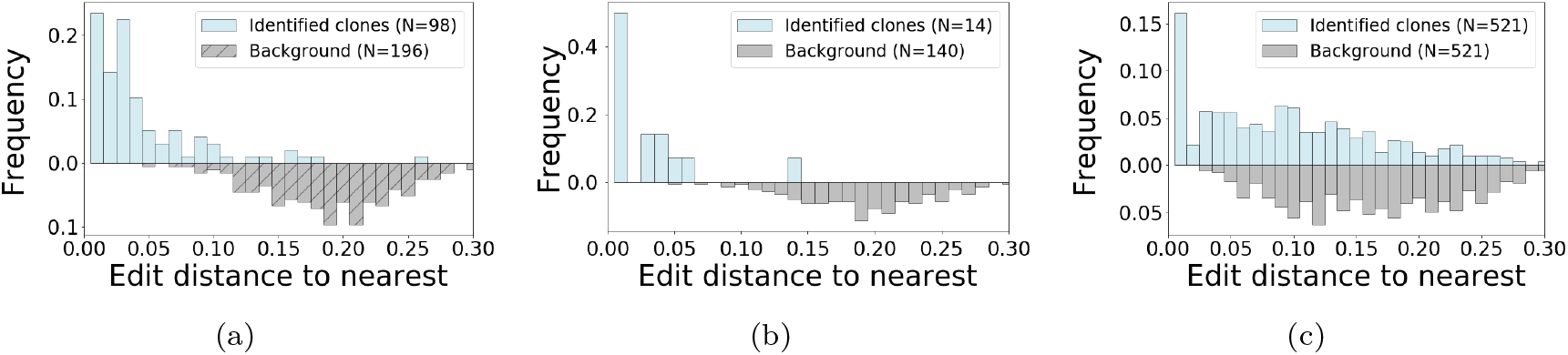
(a) Distribution of distance-to-nearest within clones with non unique J gene. (b) Distribution of distance-to-nearest within clones with non unique V gene. (c) Distribution of distance-to-nearest within clones with non unique junction length.

**Figure S4:**
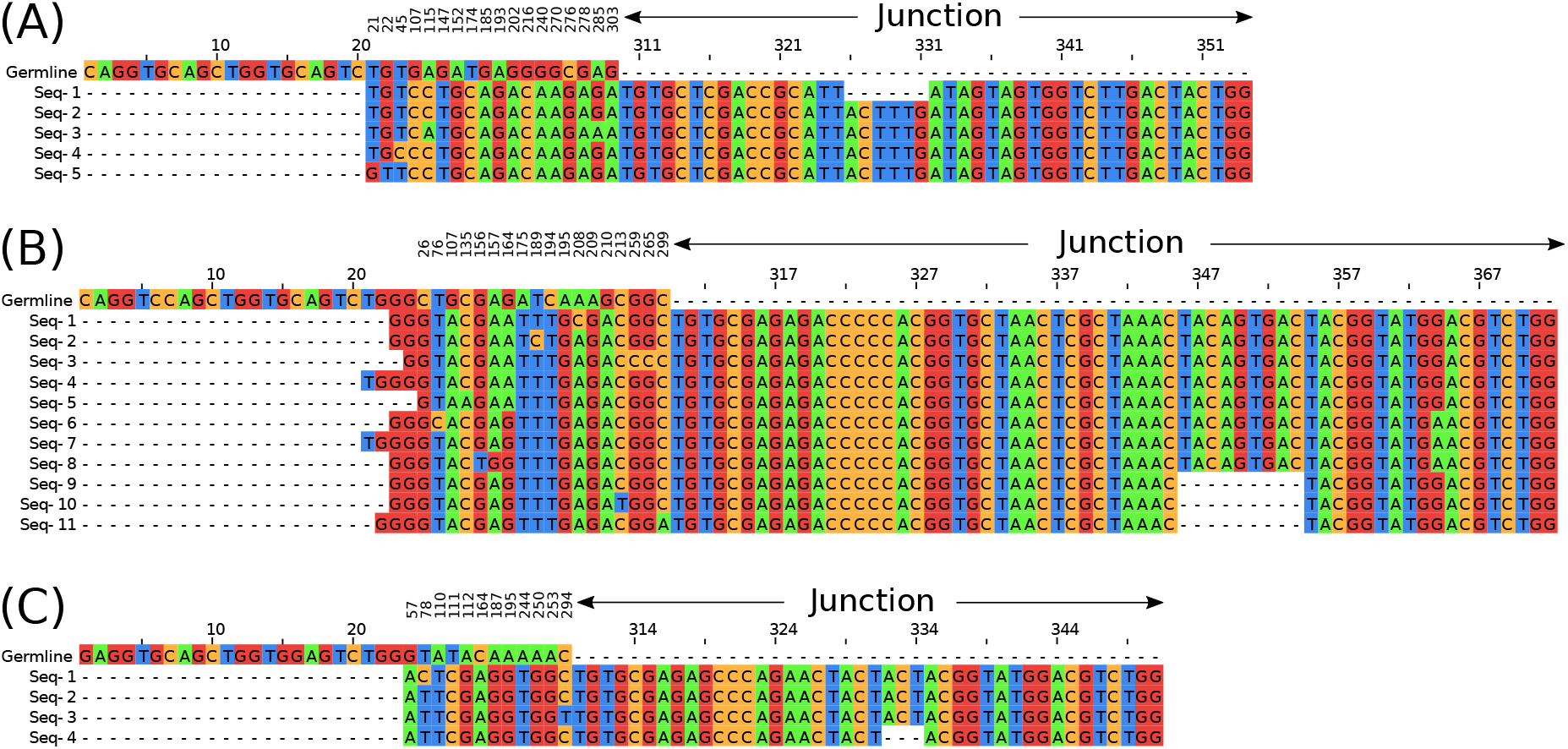
Multi-sequence alignment of the clones presented in the main text (Fig. 12). Base pairs are colored to highlight shared mutations. Top row indicates the germline, base pairs with not shared mutations are collapsed.

## Footnotes

1 https://bergvca.github.io/2017/10/14/super-fast-string-matching.html

